# Isolation and Characterization of SARS-CoV-2 strains circulating in Eastern India

**DOI:** 10.1101/2021.12.13.472526

**Authors:** Bharati Singh, Kiran Avula, Sanchari Chatterjee, Ankita Datey, Arup Ghosh, Saikat De, Supriya Suman Keshry, Soumyajit Ghosh, Amol Suryawanshi, Rupesh Dash, Shantibhusan Senapati, Tushar K. Beuria, Punit Prasad, Sunil Raghav, Rajeeb Swain, Ajay Parida, Gulam Hussain Syed, Soma Chattopadhyay

**Author notes:** Equal contributors (Bharati Singh, Kiran Avula, Sanchari Chatterjee, Ankita Datey).

## Abstract

Emergence of SARS-CoV-2 as a serious pandemic has altered the global socioeconomic dynamics. The wide prevalence, high death counts and rapid emergence of new variants urge for establishment of research infrastructure to facilitate rapid development of efficient therapeutic modalities and preventive measures. In agreement with this, five SARS-CoV2 strains (ILS01, ILS02, ILS03, ILS15 and ILS24) of four different clades (19A, 19B, 20A and 20B) were isolated from patient swab samples collected during the 1^st^ COVID-19 wave in Odisha, India. The viral isolates were adapted to *in-vitro* cultures and further characterized to identify strain specific variations in viral growth characteristics. All the five isolates showed substantial amount of virus induced CPE however ILS03 belonging to 20A clade displayed highest level of CPE. Time kinetics experiment revealed spike protein expression was evident after 16th hours post infection in all five isolates. ILS03 induced around 90% of cytotoxicity. Further, the susceptibility of various cell lines (human hepatoma cell line (Huh-7), CaCo2 cell line, HEK-293T cells, Vero, Vero-E6, BHK-21, THP-1 cell line and RAW 264.7 cells) were assessed. Surprisingly, it was found that the human monocyte cells THP-1 and murine macrophage cell line RAW 264.7 were permissive to all the SARS-CoV-2 isolates. The neutralization susceptibility of viral isolates to vaccine-induced antibodies was determined using sera from individuals vaccinated in the Government run vaccine drive in India. The micro-neutralization assay suggested that both Covaxin and Covishield vaccines were equally effective (100% neutralization) against all of the isolates. The whole genome sequencing of culture adapted viral isolates and viral genome from patient oropharyngeal swab sample suggested that repetitive passaging of SARS-CoV2 virus in Vero-E6 cells did not lead to emergence of many mutations during the adaptation in cell culture. Phylogenetic analyses revealed that the five isolates clustered to respective clades. The major goal was to isolate and adapt SARS-CoV-2 viruses in *in-vitro* cell culture with minimal modification to facilitate research activities involved in understanding the molecular virology, host-virus interactions, application of these strains for drug discovery and animal challenge models development which eventually will contribute towards the development of effective and reliable therapeutics.

## Introduction

Since its emergence in December 2019, in Wuhan, China, the Severe Acute Respiratory Syndrome Coronavirus 2 (SARS-CoV-2) has had an unprecedented effect on human health and well-being world over ^1–3^. According to WHO data, the virus has infected 240 million individuals worldwide and has so far caused 4.8 million fatalities^4^. SARS-CoV-2 is a single-stranded, positive-sense RNA virus of the *Coronavirus* genus, family *Coronaviridae* and order *Nidovirales* ^3^. SARS-CoV-2 genome is around 30 kb in size and shares 79% and 50% homology with the genome of SARS-CoV and MERS-CoV, the causative agents of two earlier coronavirus epidemics in 2002-03 and 2012. Based on the reproductive number (R_0_) SARS-CoV-2 (2-2.2) is highly infectious then SARS-CoV (1.7-1.9) & MERS-CoV (<1) ^5^.

The SARS-CoV-2 genome ORF1a/ORF1ab encodes for two polyproteins, pp1a/pp1ab which account for 2/3^rd^ of the viral genome, and the remaining 1/3^rd^ near the 3’-end encodes for four structural proteins Spike (S), Envelope (E), Membrane (M), and Nucleocapsid (N) ^1^. The overlapping pp1a and pp1ab, are proteolytically cleaved by papain-like and chymotrypsin-like viral proteases (PL^pro^ & CL^pro^) to yield 16 non-structural proteins, which play an important role in virus life cycle with cooperation from other accessory viral proteins ^6^. The transmission of this virus occurs mainly through aerosols/liquid droplets that emanate from the cough/sneeze from infected patients ^7^. Majority of the infected individuals are either asymptomatic or exhibit mild flu like symptoms, whereas few patients exhibit severe clinical manifestation leading to the severe Acute Respiratory Distress Syndrome (ARDS) ^8^.

The global prevalence of SARS-CoV-2 & rampant growth in the human host lead to emergence of mutational variability among circulating viruses. Presence of multiple variants with variability in infection/transmission and disease manifestation urge for isolation of the circulating SARS-CoV-2 variants to enhance our understanding in variant specific differences in viral growth characteristics, host interactions and disease pathogenesis. In this study, five circulating strains of SARS-CoV-2 belonging to early clades have been isolated from laboratory confirmed COVID-19 patients swab samples collected during the 1^st^ COVID-19 wave in Odisha, India. The isolated strains have been further characterized and sequenced to enable utilization of these isolates as resources in research and development towards prevention and effective therapeutic intervention against COVID-19.

## Materials and method

### Cells and Viruses

Vero E6, Vero, BHK-21, HEK293T and Huh7 cells were maintained in high glucose DMEM supplemented with 10% fetal bovine serum and 1X Pencillin/Streptomycin. CaCo2 cells were maintained in DMEM supplemented with 20% fetal bovine serum and 1X Pencillin/Streptomycin. THP-1 and RAW 264.7 were maintained in RPMI supplemented with 10% fetal bovine serum, 1X Pencillin/Streptomycin, 10 mM sodium pyruvate, 1M HEPES, and glucose. The details about all the eight cell lines are provided in Table 1. All the cell cultures were maintained in humidified environment with 5% CO_2_ at 37°C.

**Table 1:** Details of the various cell lines used in this study.

| S No | Cell line | Source |
| --- | --- | --- |
| 1 | Vero | Monkey kidney epithelial cell line |
| 2 | Vero-E6 | Monkey kidney epithelial cell line |
| 3 | HEK 293T | Human embryonic kidney cell line |
| 4 | Huh-7 | Human hepatoma cell line |
| 5 | CaCo2 | Human colon epithelial cell line |
| 6 | BHK-21 | Hamster kidney epithelial cell line |
| 7 | THP | Human monocytes cells |
| 8 | RAW 264.7 | Mouse monocytes cells |

The cells were seeded a day before infection such that they attain confluence on the day of infection. On the day of infection complete media was removed and respective virus infection was given at MOI of 0.1 in serum free media for 1.5 hr at 37°C with gentle rocking at every 15 mins. After 1.5 hr the inoculum was removed and cells were washed twice with PBS and supplemented with complete media. Five different viral strains were isolated and characterized in the current study. The details regarding these viral strains are mentioned in Table 2.

**Table 2:** Accession numbers of the genome sequence and clade information of the viral RNA from source swab samples **(S)** and isolated & culture adapted viruses **(A)** used in this study.

| Name | Accession no | Clade |
| --- | --- | --- |
| ILS01 | EPI_ISL_463010 (S) | 19A |
|  | MW559533.2 (A) | 19A |
| ILS02 | EP_ISL_3039724 (S) | 20A |
|  | EPI_ISL_1190402 (A) | 19B |
| ILS03 | EPI_ISL_463032 (S) | 20A |
|  | EPI_ISL_1196305 (A) | 20A |
| ILS15 | EPI_ISL_463054 (S) | 20B |
|  | MW828325.1 (A) | 20A |
| ILS24 | EPI_ISL_463058 (S) | 19B |
|  | MW828330.1 (A) | 19B |

### Specimen collection

Oropharyngeal swab samples collected in VTM from suspected symptomatic and asymptomatic patients by the various sample collection centres in the state of Odisha, India during April-June 2020 were used in this study. The samples were tested for presence of virus by qRT-PCR and samples with Ct (Cycle threshold) values below 15 were subsequently used for virus isolation. Upon confirmation of infection, the samples were aliquoted and kept in deep freezers until further use.

### Ethics statement

The current studies involving swab samples from the human participants were reviewed and approved by the Institutional Human Ethics Committee, Institute of Life Sciences. The Institutional Ethics Committee (IEC)/ Institutional Review Board (IRB) reference number is 96/HEC/2020. The written consent form duly signed by the participants/ legal guardian was taken into consideration for the concerned study

### Virus Isolation

Oropharyngeal swab sample**s** of confirmed COVID-19 patient**s were** used for isolation of the virus. The oropharyngeal swab sample was diluted 1:1 with DMEM supplemented with antibiotics and antifungal agents and filtered through 0.22-micron filter. Vero E6 cells were infected with the filtered swab sample for 1.5 hr at 37°C with gentle rocking every 15 mins. The inoculum was aspirated and cells washed with PBS and supplemented with fresh media containing 2% FBS. The infected cells were regularly monitored for cytopathic effect ^9^. 72 hr post infection the culture supernatants were collected and the clarified supernatant (at 3000 rpm for 5 mins) were used as inoculum for subsequent (2^nd^) passage of virus in naїve Vero E6 cells. This process was repeated every 48 hrs up to the 10^th^ passage. RNA isolated from the culture supernatants was used for confirmation of SARS-CoV-2 virus isolation by qRT-PCR ^10^. Virus titres in the culture supernatants was estimated by TCID_50_ assay. RNA isolated from 10^th^ passage virus was used for determining the whole genome sequence. SARS-CoV-2 virus isolation and culture was conducted in the biosafety level-3 containment facility according to the guidelines issued by the Department of Biotechnology, Government of India. This study has been approved by the Institutional biosafety committee (IBSC) (IBSC file no. V-122-MISC/2007-08/01).

### Viral RNA extraction & estimation

RNA isolation from culture supernatant was performed using QIAamp Viral RNA Kit (Qiagen, cat. no. 52906) according to the manufacturer’s instructions. The isolated RNA was subjected to qRT-PCR for determining the viral load by absolute quantification by real-time RT-PCR using Takara PrimeScript™ one-step RT-PCR Kit (RR055A) with forward (5′-GTGAAATGGTCATGTGTGGCGG-3′) and reverse (5′-CAGATGTTAAAGACACTATTAGCATA-3′) primers and probe (5′-FAM-CAGGTGGAACCTCATCAG GAGATGC-BHQ-3′) targeting the SARS-CoV2 RdRp gene. Standard curve was generated using known quantities of SARS-CoV2 viral RNA purified from the viral stock supernatants.

### Plaque Assay

To determine the viral titre plaque assay was performed as described by Mishra et.al (2016) ^11^. In brief, 80% confluent VeroE6 cells were infected with serially diluted viral culture supernatant. Subsequently the cells were overlaid with complete methyl cellulose and maintained in the incubator at 37ºC with 5% CO2. After the development of the visible plaques (6-7 days), the plaques were fixed by adding 8% formaldehyde. Later on, the cells were stained using crystal violet. The number of plaques were counted as plaque forming unit/mL (PFU/mL).

### TCID50 Assay

Vero E6 cells seeded at 90% confluency in 96-well plates were infected for 1 hr at 37°C with 100 uL of serially diluted (10-fold) virus inoculum in DMEM with 2% FBS. 1 hr post infection the inoculum was aspirated and cells were replenished with fresh media. 3 days post infection the cells were fixed in 4% paraformaldehyde and stained with 1% crystal violet to determine the cytopathic effect. Median tissue culture infectious dose (TCID50) was determined by the Reed and Muench method ^12^.

### Immunofluorescence Assay

The immunofluorescence assay was performed according to the method described by Kim et.al (2013) for the detection of infected cells ^13^. The Vero E6 cells grown on glass cover slips were infected with 0.1 MOI of respective isolates and 48 hr post infection fixed in 4% paraformaldehyde. Subsequently the cells were permeabilized and blocked for 1 hr with PBS containing 0.1% TritonX-100 and 3% BSA, followed by incubation with antibody targeting the SARS-CoV-2 nucleocapsid (Abgenex, cat. No. 11-2003) overnight at 4ºC. After 3x washes with PBS, the cells were stained with the respective Alexa Fluor conjugated secondary antibody (Invitrogen, Carlsbad, CA), for 1 hr at room temperature followed by 3x washes with PBS. After the final wash, the coverslips were mounted onto ProLong Gold Antifade (Invitrogen, Carlsbad, CA). Images were captured under a 100×oil immersion objective lens using a Leica TCS SP5 Confocal microscope for detection of virus-infected cells protein.

### Western blot analysis

Immunoblot analysis was carried out as mentioned before^13^. In brief, cells were lysed in RIPA buffer (20 mM Tris-HCl [pH 7.5], 150 mM NaCl, 50 mM NaF, 1 mM Na3VO4, 0.1% SDS, and 0.5% TritonX-100) containing the protease inhibitor cocktail (Thermo Scientific). The whole cell lysates (WCL) were subjected to SDS-PAGE and transferred to nitrocellulose membrane (Thermo Scientific) followed by blocking and immunoblotting with antibodies specific for SARS-CoV-2 spike (Abgenex, cat. No. 10-1007) and nucleocapsid (Abgenex, cat. No. 11-2003).

### Micro-neutralization Assay

Micro neutralization assay was performed as mentioned before ^14^. Briefly, serum samples were heat-inactivated for 60 minutes at 56°C; and syringe filtered through 0.22 µm. These samples were then two-fold serially diluted in a 96-well plate starting from 1:10 and then mixed with equal volume of virus solution containing 1000 TCID_50_ of SARS-CoV-2. This serum-virus complex was incubated for 1 hour at 37°C followed by addition in duplicate to a 96 well plate containing 90% confluent Vero E6 monolayer. The plates were incubated for 36 hours at 37°C in a humidified atmosphere with 5% CO2 (Ref 6). Afterwards, the cells were washed and fixed with 4% paraformaldehyde followed by blocking with 2% BSA for 1 hr at room temperature. Cells were then incubated with SARS-CoV-2 rabbit anti-nucleocapsid (Abgenex, cat. No. 11-2003) antibody for 1-2 hour followed by 3x wash with PBS and 1 hr incubation with horseradish peroxidase-conjugated goat anti-rabbit IgG. After 3x wash, equal volume of 3,3’,5,5’-tetramethylbenzidine substrate was added to each well for 15 minutes with termination of reaction by addition of 2N H_2_SO_4_. The plates were read at 450/620 nm using a microplate reader. The neutralization percentage was determined by following the formula: each well is 100 – [(X-average of ‘no virus’ wells)/ (average of ‘virus only’ wells - average of ‘no virus’ wells) *100], where X is the read for each well. Non-linear regression curve fit analysis over the dilution curve was performed in the Graphpad Prism 5 software while setting the top and bottom constraints at 100% and 0% ^15,16^.

### Viral Genome sequencing and analysis

For the whole genome sequencing of the isolated viruses, the viral RNA amplicon libraries were prepared using the QIAseq FX DNA Library Kit and the QIAseq SARS-Co V-2 Primer Panel (Qiagen, cat. no. 180475, cat. no. 333896) as instructed by the manufacturer’s manual. The library was sequenced using the Illumina platform. The adapter sequence used for each sample was compatible with the Illumina NextSeq 550 instrument with 96-sample configurations (Qiaseq unique dual Y-adapter kit). The average insert length was in the 250–650 bp range. The raw data pre-processing, alignment with viral genome, consensus sequence generation, variant calling and phylogenetic analysis was performed as described by Raghav et al., 2020 ^17^.

### Statistical Analysis

Statistical analysis was performed using the GraphPad Prism software version 5. Data were presented as mean ± standard deviation (SD). The Non-Linear fit log (inhibitor) vs. response - Variable slope was used to determine the percentage inhibition of virus infection due to vaccine-induced antibody-mediated neutralization.

## Results

There is an urgent need to isolate and establish culture of the SARS-CoV-2 circulating viral strains to aide in research and development towards finding efficient therapeutic modalities and vaccine development. Hence attempts were made to isolate SARS-CoV-2 virus from COVID-19 patients oropharyngeal swab samples collected during April-June 2020 at various location in the state of Odisha, India. The viral RNA obtained from the swab samples was subjected to whole genome sequencing to identify the viral strain and emerging mutations. Based on the whole genome sequencing result and cycle threshold values of qRT-PCR swab samples were chosen, which were expected to have high viral load and of respective clades 19A, 19B, 20A & 20B for virus isolation and propagation.

Virus isolation was carried out using the protocol adapted by Harcourt et al, 2020 with minor modification. Based on previous reports Vero-E6 cells were used for virus propagation ^9^. All the five isolates were passaged for ten times on Vero-E6 cells and the culture supernatants were collected during every passage and a portion of clarified supernatant was used as inoculum for subsequent passage. The 10^th^ passage clarified supernatant was used as viral stock for the whole genome sequencing, virus characterization, and further experiments. During the passages, RNA isolated from the collected supernatants was subjected to qRT-PCR to confirm the presence of SARS-CoV-2. Viral titres in the 10^th^ passage supernatant were determined by standard plaque (**Figure 1A**) and TCID_50_ assays (**Figure 1B**). The viral titres of the respective isolates ranged from the 10^6^ to 10^8^/mL. Based on the titres obtained, the Vero-E6 cells were infected with 0.1 MOI of all isolates for subsequent experiments. To visualize the cytopathic effect (CPE) bright field images were captured at 48 hr post infection. All the five isolates displayed significant amount of virus induced CPE however ILS03 belonging to 20A clade displayed highest level of CPE among the five and the other four displayed nearly similar levels of CPE (**Figure 1C**). Absolute quantification of viral genome copies in the culture supernatant collected at 48 hr post infection was determined by qRT-PCR using gene specific primers and probes for nucleocapsid and ORF1 (**Figure 1D**). Viral gene expression was also confirmed by Western blot analysis of the cell lysates using antibodies targeting SARS-CoV-2 spike and nucleocapsid (Figure 1E). Cells infected with all 5 isolates showed profound level of viral gene expression as adjudged by Western blot analysis. To determine the level of infectivity or any isolate specific variation in subcellular infection pattern immunofluorescence assay was performed in Vero-E6 cells infected with the respective isolates at 0.1 MOI for 48 hours. No significant variation was observed in the subcellular distribution of the SARS-CoV2 nucleocapsid protein and all the isolates displayed reticular cytoplasmic staining across the entire cytoplasm (**Figure 1 F**). Quantification of the percentage of infected cells showed that around 70-90% of cells were infected at 48 hours post infection with the respective isolates using 0.1 MOI (**Figure 1 G**).

**Figure 1:**
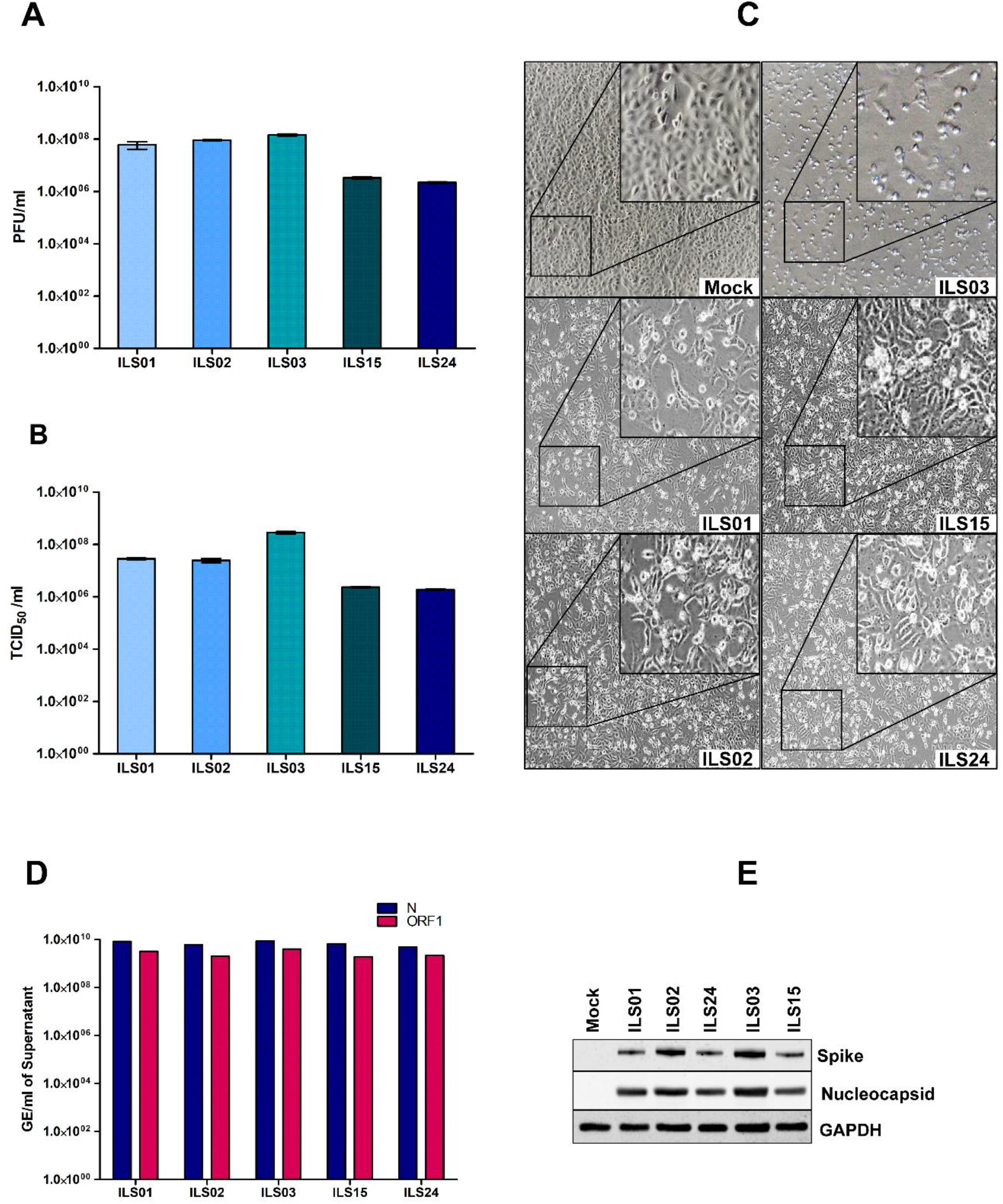
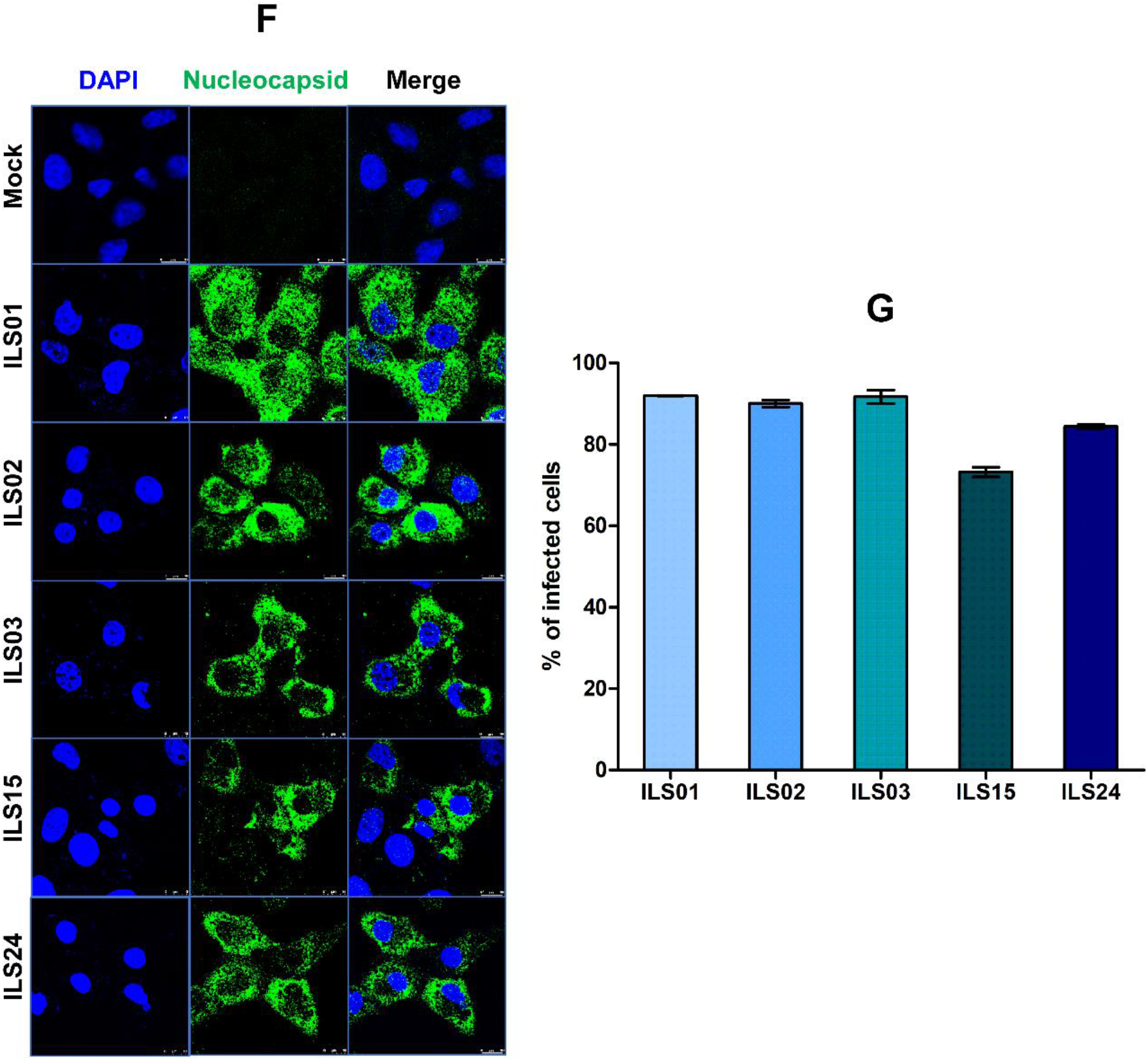
Characterization of isolated SARS-CoV2 circulating strains. The SARS-CoV2 circulating strains were isolated from the COVID-19 patients swab samples through sequential passage in Vero E6 cells as described in materials & methods. The viral titres, cytopathic effect and gene expression was determined in the 10^th^ passage viral stocks. Quantification of viral titres of the five isolates by plaque forming unit (PFU) assay **(A)** and TCID_50_ assay **(B)**. Bright field images depicting cytopathic effect in Vero E6 cells infected respectively with the five isolates **(C)**. Absolute quantification of viral genome copies in all five isolates using gene-specific primer and probes targeting SARS-CoV2 nucleocapsid and ORF-1 gene **(D)**. Western blot analysis of infected Vero E6 cell lysates with antibodies against SARS-CoV2 spike and nucleocapsid **(E)**. GAPDH was used as protein loading control. Immunofluorescence detection of SARS-CoV-2 infected cells using antibody against SARS-CoV2 nucleocapsid in Vero E6 cells infected with 0.1 MOI of respective isolates **(F)** and quantification of the percentage of infection 48 hr post infection **(G)**.

To access the relative differences in the kinetics of viral gene expression, time kinetics experiment was conducted by infecting Vero-E6 cells at MOI of 0.1 and collecting cells at every 4 hours interval for 24 hrs. Western blot analysis of the cell lysates for spike and nuclocapsid proteins of SARS-CoV-2 showed that in isolates ILS01, ILS02, & ILS03, nucleocapsid expression is noticeable from the 16th hour post infection, whereas in isolates ILS15 and ILS24, it appears from 12th hour onwards (**Figure 2**). Interestingly, spike protein expression was evident only after 16th hours post infection in all five isolates (**Figure 2**). To further access the specific variations between the isolates in virus mediated cytotoxicity and viral replication kinetics, Vero E6 cells were infected at 0.1 MOI with respective isolates and the cell culture supernatants collected every 12 hours upto 60 hours post infection to estimate cytotoxicity and viral release. Based on the LDH levels in the supernatants, it appears that isolates ILS01, ILS02, ILS15 & ILS24 induce around 30% of cytotoxicity with respect to mock at 48 hr post infection, whereas isolate ILS03 induces around 90% of cytotoxicity (**Figure 3A-3E**). Quantification of viral genome copies in the culture supernatants suggest a steady increase in the genome copies from 12 to 36 hr post infection indicating that there is an exponential increase in the release of viral particle upto 36 hrs post infection followed by plateau (**Figure 3F**).

**Figure 2:**
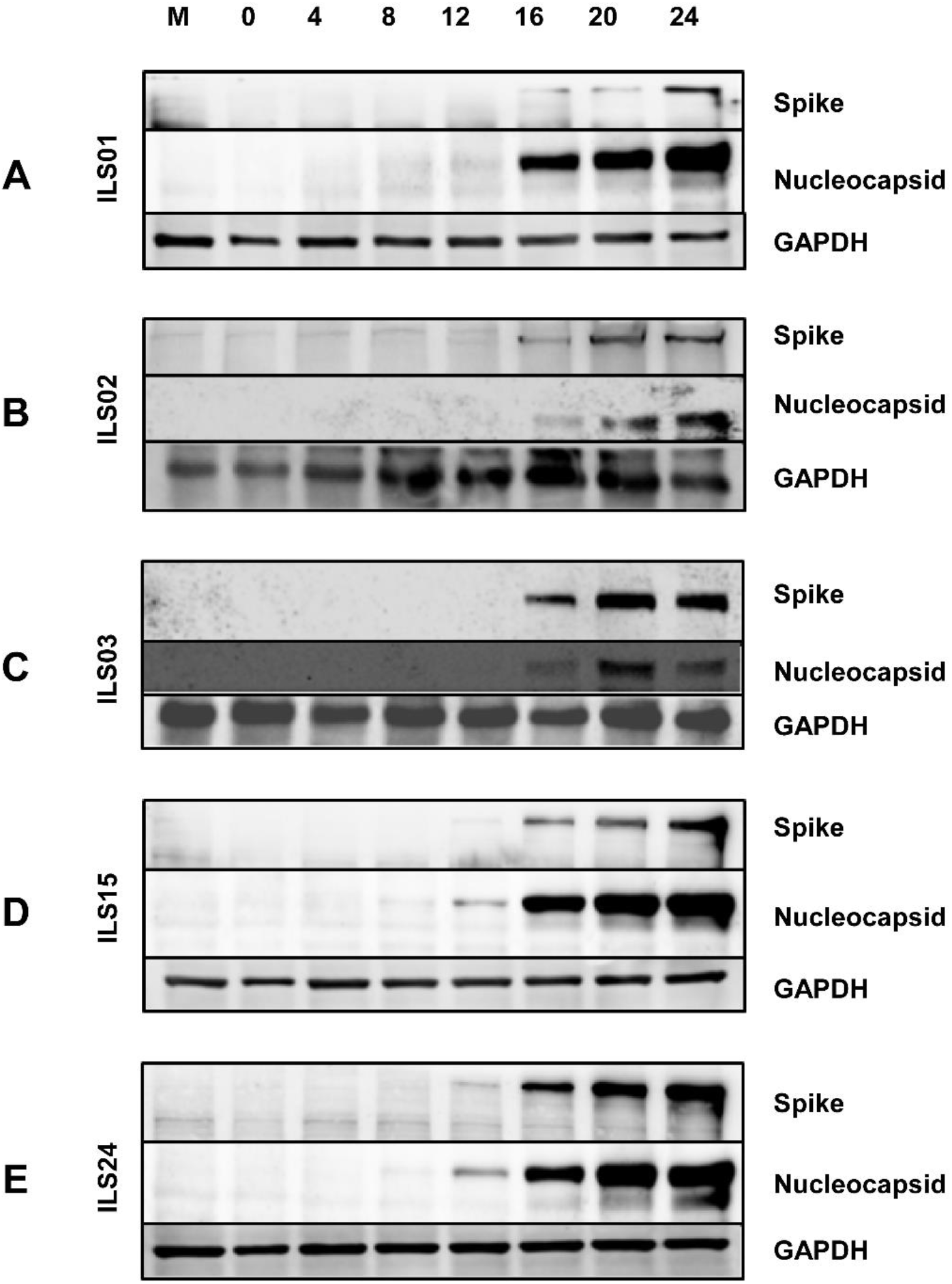
Time kinetics of viral gene(s) expression. Vero E6 cells infected with respective isolates of SARS-CoV2 were collected at indicated time points post infection. Cell lysates were subjected to Western blot analysis with antibodies against SARS-CoV2 spike and nucleocapsid proteins. GAPDH was used as an internal loading control.

**Figure 3:**
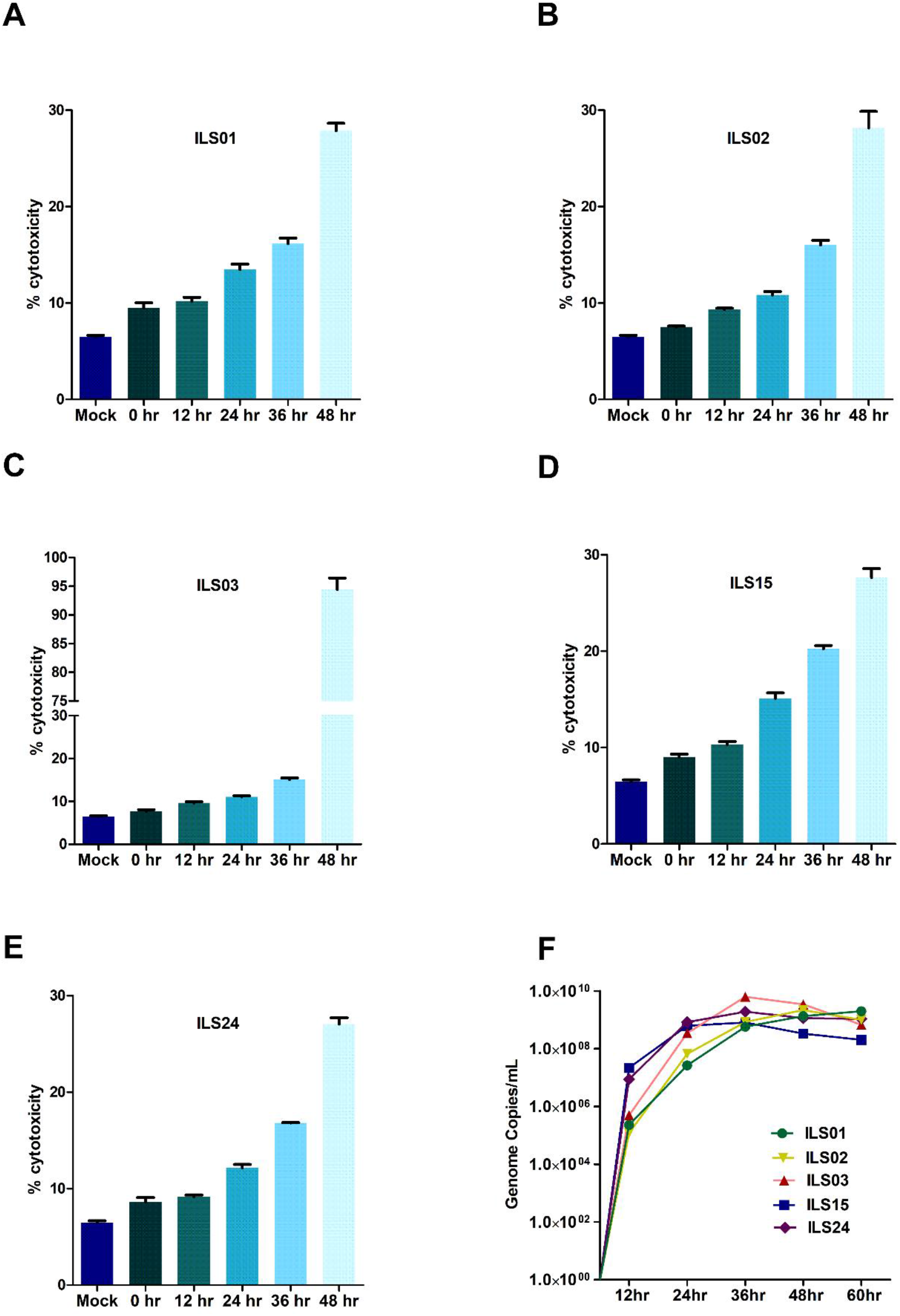
Viral cytopathy and growth kinetics. Infection associated cytopathy was determined by calculating LDH release as described in materials & methods. **(A-E)** Graph depicting percentage of cytotoxicity in the infected Vero E6 cells at respective time points post infection. **(F)** Line plot showing time-dependent increase in the viral genome copies in culture supernatants determined by absolute quantification of viral genome.

Further the susceptibility of various cell lines was assessed towards isolates to decipher isolate specific variations in cell susceptibility. Various cell lines were infected with the respective isolates at 0.1 MOI and culture supernatants were collected at 24 hrs post infection (hpi) to quantify the viral genome copies. The human hepatoma cell line (Huh-7), which is highly susceptible to Dengue, Chikungunya, and Hepatitis C viruses (HCV), was found to be more or less equally susceptible to all the five isolates (**Figure 4A**). Similarly, CaCo2 cell line, which is a human intestinal epithelial cell line that has been shown by various groups to be permissive to SARS-CoV-2 was also found to be susceptible to all the five isolates (**Figure 4B)**. However, the isolate ILS01 was found to be less infectious compared to the other isolates. Similarly, HEK-293T cells (a human kidney cell line) was found to be more permissive to isolate ILS01, ILS02, ILS15 & ILS 24 as compared to isolate ILS03 (**Figure 4C)**. Immune cells predominantly show selective susceptibility to the viruses. Surprisingly, in our study, we found that the human monocyte cells THP-1 and murine macrophage cell line RAW 264.7 were permissive to all the SARS-CoV-2 isolates (**Figure 4D and 4E**).

**Figure 4:**
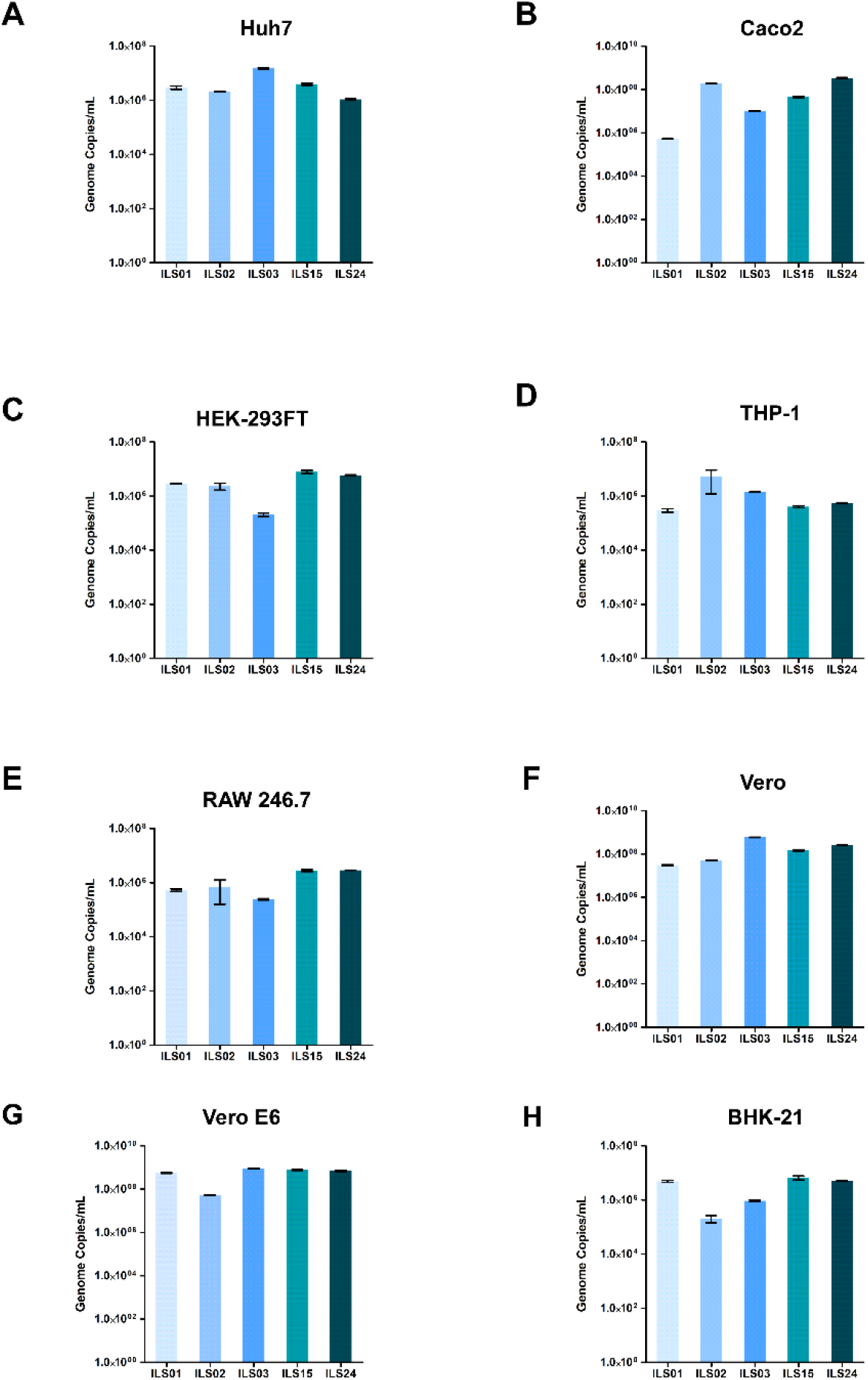
Susceptibility of various cell lines to the SARS-CoV2 isolates. Different cell lines were subjected to infection with 0.1 MOI of respective isolates. 24h post infection the viral load in the culture supernatants was determined by absolute quantification of viral genome copies. Graphs depicting the viral copies per ml supernatant in Huh7 **(A)**, Caco2 **(B)**, HEK 293T **(C)**, THP1 **(D)**, RAW 264.7 **(E)**, Vero **(F)**, Vero E6 **(G)**, and BHK-21 **(H)**.

To decipher any clade specific variations towards neutralization, the neutralization capacity and protection of the vaccine-induced antibodies against the respective isolates was determined. Neutralizing antibody levels predict vaccine efficacy and immune protection. In India, initially only two vaccines, Covaxin and Covishield were given emergency approval and used in Government run COVID-19 vaccination drive. We used vaccinated sera from Covaxin and Covishield vaccinated healthy individuals with no history of SARS-CoV-2 infection. The sera were collected after completion of the 2^nd^ vaccine dose fifteen days post 2^nd^ vaccine dose. Horse sera was used as negative control as it was difficult to obtain age-matched healthy control sera from individuals who had not been vaccinated or exposed to COVID-19. The micro-neutralization assay suggested that both the vaccine was equally effective against all of the isolates. Nearly 100% neutralization was observed at 1:10 dilution, which declined to ∼50% at dilutions 1:160 or higher **(Figure 5)**.

**Figure 5:**
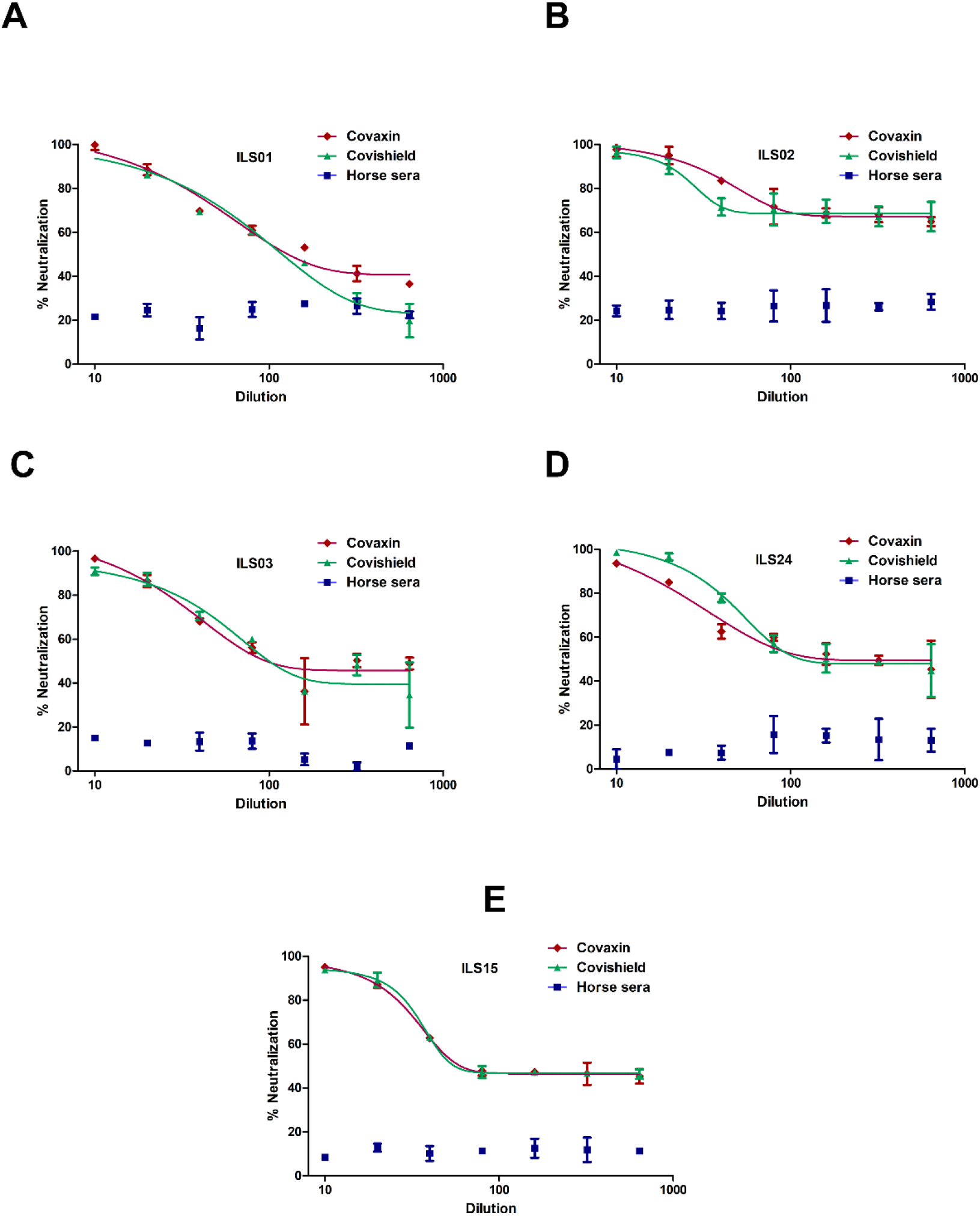
Neutralization potential of sera obtained from vaccinated individuals:. The respective isolates were subjected to micro-neutralization assay using the sera obtained from Covaxin and Covishield vaccinated individuals to determine the neutralization potential of the post vaccination sera against the respective isolates. The dose-response curves were fitted using a nonlinear regression model using the GraphPad software Prism 5. **(A-E)** Neutralization efficiency of the respective vaccinated sera against the 5 isolates. Horse sera was used as negative control.

The whole genome sequencing of culture adapted viral isolates and viral genome from patient oropharyngeal swab sample suggested that repetitive passaging of SARS-CoV2 virus in Vero-E6 cells did not lead to emergence of many mutations during the adaptation in cell culture. The number of viral gene mutations found in the source swab samples and isolated viruses in comparison to the Wuhan reference strain is shown in **Table 3**. Comparative analysis of common and unique sequence mutation between the source sample and isolate (**Table 4)** and mutational plot analysis of non-synonymous mutations **(Figure 6)** suggests that during the culture adaptation very minimal changes occurred. ILS01 isolated from source sample of clade19A gained only one mutation (A23014C) in spike gene during cell culture adaptation, while it retained all other ten mutations found in source swab samples. Isolate ILS24 obtained from source samples of clade 19B gained three mutations (C2143T, C10138T, C10702T) in the ORF1ab and one mutation (G28326T) in the N genes during adaptation. It retained 5 mutations found in the source swab sample material, and one reversion (G26730T) to Wuhan reference strain in the M gene. ILS03 isolated from swab sample of clade 20A retained 9 mutations found in swab sample and gained one mutation each in ORF1ab (G19514T) and S (A24538C) genes during adaptation. Interestingly during isolation and adaptation of ILS15 from swab sample of clade 20B, five reversions occurred, which included two (C8917T, G9389A) in ORF1ab and three (G28882A, G28881A, G28883C) in N gene resulting in the reclassification of the cell culture adapted strain ILS15 in clade 20A. To understand the evolution of the virus and trace lineage phylogenetic network analysis was performed using the genome sequence of the four isolates and 33 other largely complete sequences of SARS-CoV-2 genome from different regions of the world. Phylogenetic analysis indicated that the genome sequence of the swab sample and culture adapted viruses remain identical as they cluster close together in the respective clades (**Figure 7**), which was also in agreement with the mutational plot analysis evident by the presence of similar nonsynonymous mutation throughout respective genomes (**Figure 6**). Both the swab sample and adapted virus of isolates ILS01& ILS24 closely clustered together with the Wuhan reference strain as they belong to very early clade 19A & 19B respectively. Swab sample in case of isolate ILS15 cluster together with viral genome from India & Brazil belonging to clade 20B whereas the adapted virus strain cluster together with the genome sequences from Australia and South Korea of clade 20A which may be due the 5 reversions found in the adapted virus. Interestingly, in case of isolate ILS03 both the swab sample and adapted virus strain extended out and clustered separately from the other viral genome used in this analysis.

**Table 3:** Tabular representation of genome sequences of all 5 isolates with reference to Wuhan strain (NC_045512).

| Sample | Name | Clade | Total Coverage | Missing region | # of missing bases | # Of nonsynonymous mutation |
| --- | --- | --- | --- | --- | --- | --- |
| Swab (S) | ILS01 | 19A | 29665 | 1-30 | 30 | 11 |
|  | ILS02 | 20A | 29836 | 1-54,521-530 | 64 | 27 |
|  | ILS03 | 20A | 29688 | 1-2 | 2 | 9 |
|  | ILS15 | 20B | 29680 | 1-33 | 33 | 9 |
|  | ILS24 | 19B | 29805 | 1-30 | 30 | 6 |
| Adapted Virus (A) | ILS01 | 19A | 29836 | 1-32 | 32 | 12 |
|  | ILS02 | 19B | 29873 | 1-4 | 4 | 11 |
|  | ILS03 | 20A | 29836 | 1-4 | 4 | 11 |
|  | ILS15 | 20A | 29836 | 1-29 | 29 | 6 |
|  | ILS24 | 19B | 29836 | 1-29 | 29 | 9 |

**Table 4:**
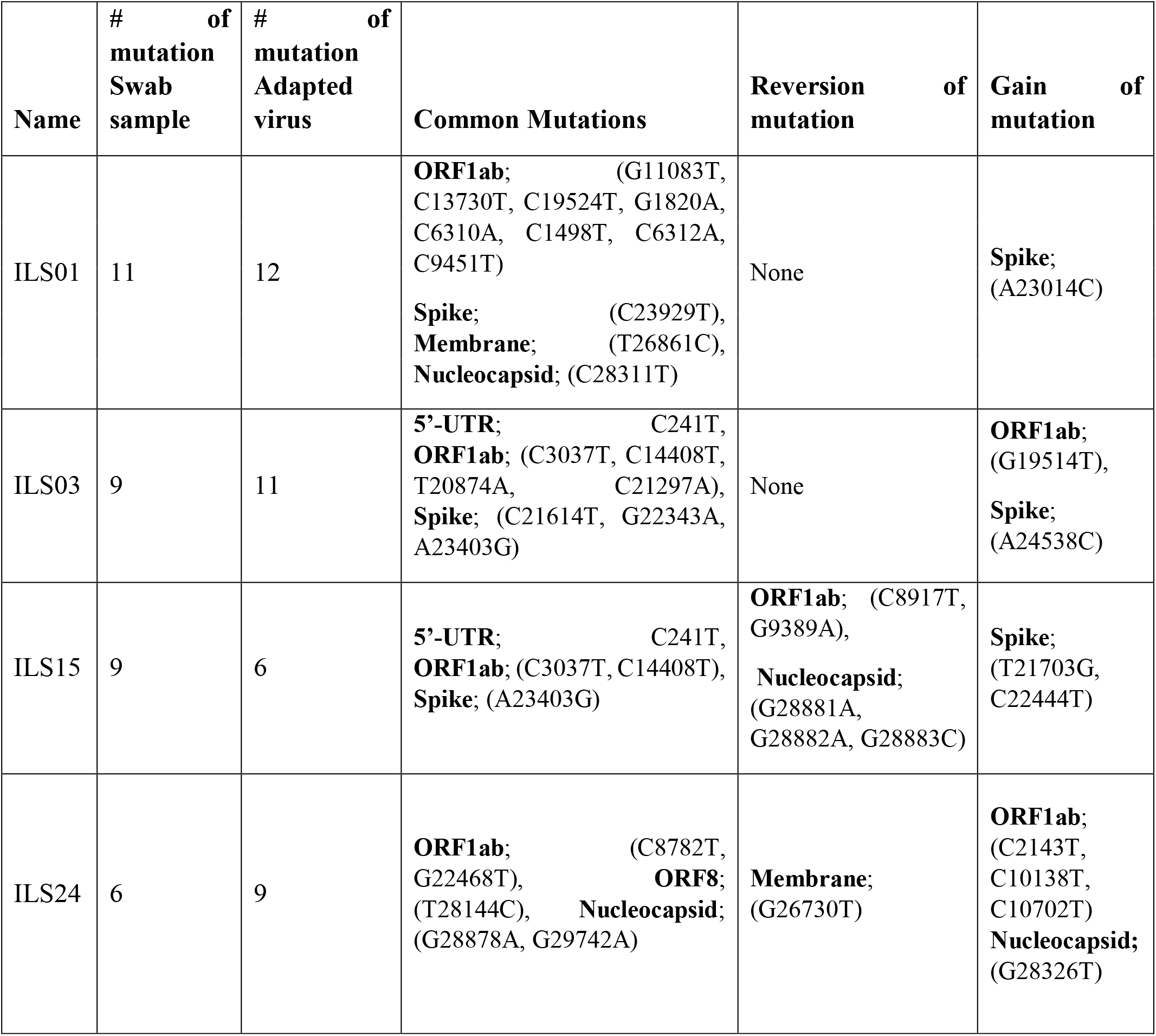
Tabular representation of SARS-CoV2 gene-specific non-synonymous mutations in both the swab samples and cell culture adapted strains.

**Figure 6:**
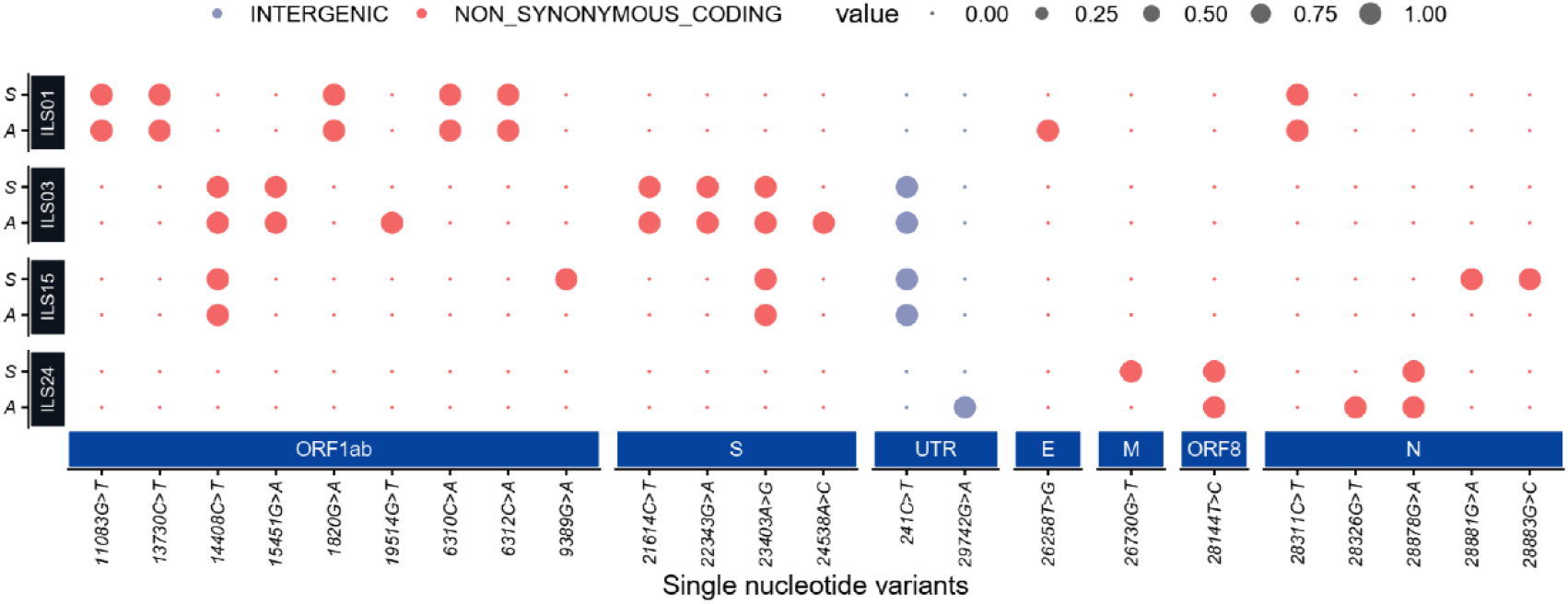
Mutation plot of the isolates and source swab samples:. Dot plot representing high quality single nucleotide nonsynonymous and intergenic variant (SNV) present in the initial viral RNA isolated from patients swab samples (denoted as **S**) and viral RNA from culture adapted isolates (denoted as **A**). The large dot represents the presence of a SNV in the represented sample coloured by their functional annotations (grey for intergenic, red for non-synonymous SNVs).

**Figure 7:**
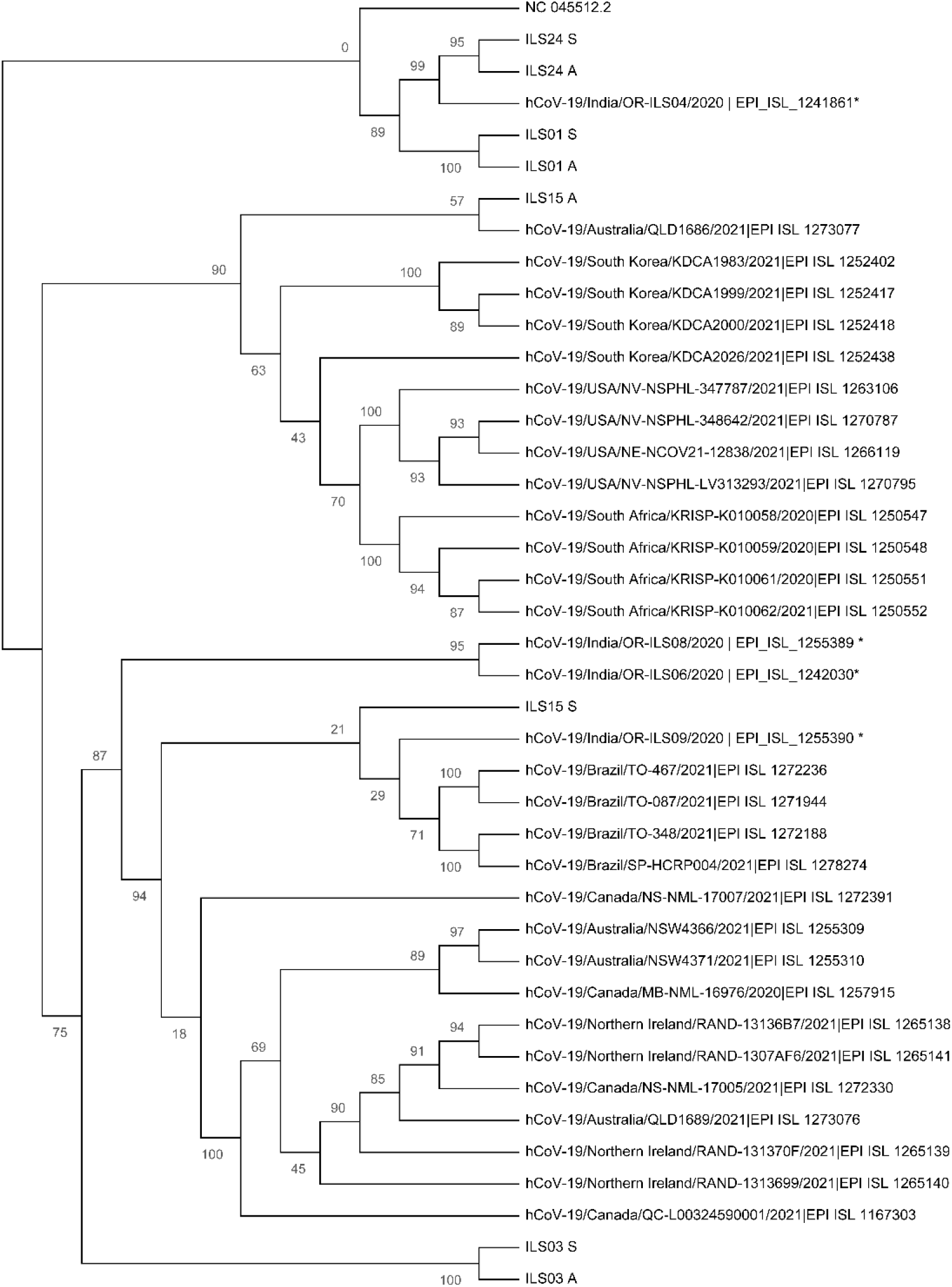
Phylogenetic network analysis of the isolated viruses:. Maximum likelihood (ML) tree of studied viral sequences in combination with 33 SARS-CoV2 genome sequences representing from different countries around the globe including four sequences from Odisha India. Bootstrap (n =1000) values are represented as branch labels.

## Discussion

In the prevailing pandemic state, it is important to isolate and characterize the disease-causing pathogen to facilitate development of therapeutic strategies and vaccine candidates. Therefore, in this study we have isolated and characterized five circulating local strains of SARS-CoV-2 as limited COVID-19 resources were available in India to aide in research and development.

As done by other groups Vero-E6 cells were used for the isolation of SARS-CoV-2 viruses ^18,19^. We observed a robust virus-induced cytopathic effect from 5^th^ passage onwards similar to previous reports ^20^. The viral titres were around 1×10^6^ TCID _50_ /ml in final passages for all the isolates **(Figure 1A & 1B)** similar to the titres reported by other groups ^21,22^. Subsequent infection with the isolated viruses leads to robust infection in Vero-E6 cells, which was evident by exponential increase in virus release from 12-36 hours post infection and detection of infection in 80-100% of Vero-E6 cells, 48 hours post infection. In agreement with studies from other labs the isolates of the current study also showed infectivity in various cell lines ranging from primate to human epithelial & immune cells. The immune cells have been shown to display selective susceptibility to some viruses. For example the THP-1 monocyte cells are not permissive to HCV and Chikungunya viruses ^23,24^, whereas they permissive to Dengue virus ^25^. In this study it was found that the viral replication levels of all the isolates were nearly similarly in immune cells in comparison to the cells of epithelial lineage. Although the viral growth kinetics was similar between ILS03 and other isolates, ILS03 displayed 2 fold higher cytopathic effect compared to other isolates suggesting that the high CPE observed with ILS03 might be due to unique characteristics of ILS03 and not due to mere high viral load **(Figure 3)**. However, further studies are warranted to characterize mechanism specific to ILS03-mediated CPE and decipher isolate-specific variations in host-virus interactions. Our current observations suggest that all the five isolates belonging to the four different clades showed almost similar virus growth characteristics despite the genomic variations between the clades suggesting that the adaptive evolution occurring in the natural host may not be applicable to growth *in-vitro* in cells highly permissive to viral infections.

In natural environment SARS-CoV-2 evolves at an estimated nucleotide substitution rate ranging between 10^−3^ and 10^−4^ substitutions per site per year ^26^ which is a very slow mutational rate. However, the rapid emergence of SARS-CoV2 variants has been speculated to have happened in chronically infected immunosuppressed patients with high levels of viral replication for extended periods under conditions of challenge with treatment modalities like transfusion of convalescent plasma or broadly neutralizing monoclonal ^27^ driving the selection of variants that evade antibody responses. However, the high prevalence of SARS-CoV2 during the past years and the rampant growth in the human host may have also contributed to mutational variability among circulating viruses. In natural host due to higher barrier towards infection, the viruses evolve and variants with higher replicative fitness get selected over time, however in *in vitro* cell cultures using highly permissive cell lines the barrier against viral replication is very low which may not favour rapid evolution of viral variants. In correlation, minimal number of mutations were observed in the adapted viruses as compared to their source swab sample even after 10^th^ passage **(Table 4, Figure 6)** suggesting that *in-vitro* cultured viruses are highly stable.

The five isolates used in this study belong to the four clades (19A, 19B, 20A, & 20B) with the clades 20A & B harbouring the D614G mutation in spike protein which has been suggested to promote higher infectivity and transmission ^17^. The observations of the current study suggest that the two vaccines, Covaxin and Covishield are equally effective and offer protection against these viral isolates from samples collected during the 1^st^ wave of COVID-19 in Odisha, India. The Covaxin is a whole inactivated virus (strain NIV 2020-770) and Covishield (Chimpanzee Adenovirus encoding the SARS-CoV-2 spike glycoprotein (ChAdOx1-S) based on the early viral isolates closer to the Wuhan strain. However, during the 2^nd^ wave many new variants were emerged across the world and they escaped neutralization by antibodies induced by vaccines based on early isolates. Majority of the neutralizing antibodies found in convalescent sera target the spike and RBD domain of spike ^28,29^, therefore many organizations have adapted the strategy of developing vaccine candidates based on Spike protein. However, further studies are warranted to evaluate the efficacy of vaccines based on whole inactivated viruses and other the antigenic motifs other than spike as they can induce a broad antibody response that may be effective against the spike variants. Use of vaccine cocktails may also be an effective strategy to overcome the burden of vaccine escaping viral variants. In agreement, recent evidence suggests that heterologous prime-boost vaccination strategy is more effective alternative than homologous prime-boost vaccination strategy against the emerging variants ^30^.

In summary, in the current investigation virus cultures of five SARS-CoV-2 strains belonging to various clades were established from the laboratory-confirmed SARS-CoV-2-infected patients and their growth kinetics and genome sequences were characterized. Further studies are required to clearly elucidate the strain specific variation among the isolates. These isolates will be highly useful resource to facilitate research and development in the field of coronavirus biology and COVID-19.

## Acknowledgement

The authors acknowledge the support of the Institute of Life sciences in conducting these studies and acknowledge the support staff involved in BSL3 maintenance, swab sample collection and processing. The authors acknowledge the financial support from DBT-ILS. SaC acknowledges CSIR for her fellowship. GHS acknowledges the Intermediate Fellowship from the DBT-Wellcome Trust India Alliance (IA/I/15/1/501826).

